# Minimal Correlation but Complementary Diagnostic Utility for Plasma Cell-free RNA and Proteins

**DOI:** 10.1101/2025.04.09.647972

**Authors:** Andrew Bliss, Conor J. Loy, Jihoon Kim, Chisato Shimizu, Joan S. Lenz, Emma Belcher, Adriana H. Tremoulet, Jane C. Burns, Iwijn De Vlaminck

## Abstract

Proteins and RNA circulate in plasma and can offer insights into human physiology. Yet, despite their clinical importance, direct comparisons between these analytes remain unexplored. Here, we measured and compared plasma cell-free RNA (cfRNA) and protein levels for 263 children diagnosed with inflammatory diseases by RNA-sequencing (n=155) and SomaScan proteomics (n=171). Remarkably, cfRNA and protein levels were largely uncorrelated across samples (feature-by-sample r=0.052; median feature-level r=0.009). Nonetheless, machine learning models based on either modality distinguished Kawasaki Disease (KD) from Multisystem Inflammatory Syndrome in Children (MIS-C) with similar high accuracy (median AUC > 0.93). Analysis of KD subtypes revealed distinct cfRNA and protein signatures, with one group showing molecular similarity to MIS-C. These findings underscore the complementary nature of cfRNA and protein profiling and highlight the utility of integrating multiple blood analytes to improve disease classification and deepen our understanding of complex inflammatory conditions.

## INTRODUCTION

Blood-borne bio-analytes such as cfRNA and proteins in plasma are increasingly used in diagnostic medicine and therapeutic development. Protein biomarkers are frequently used in translational research and clinical practice due to their biological origin, their direct or indirect roles as therapeutic targets, and their accessibility in plasma. Recent advances in high-throughput proteomics, including mass spectrometry and affinity-based assays, have expanded the quantifiable proteome to thousands of distinct proteins^1^. Despite this promise, high-throughput proteomics assays still face challenges. Mass spectrometry often requires extensive pre-fractionation and complicated sample preparation, which can limit throughput, increase costs, and introduce variability in large-scale analyses^1,2^. Affinity-based assays, including Olink and SomaScan, circumvent some of these issues but are constrained by antibody or aptamer specificity and remain expensive, limiting their widespread use in routine diagnostics and drug screening^1^. Recent comparisons of SomaScan and Olink have shown limited agreement^3^ (median rho = 0.26), drawing into question the best platform for plasma-based proteomics. Moreover, the large dynamic range of protein concentrations in plasma adds further complexity for detection and quantification irrespective of the assay platform^4^.

cfRNA is an emerging circulating bio-analyte that may address some of these challenges. cfRNA can be measured by RNA sequencing which tends to be highly reproducible and benefits from a continued reduction in cost^5^. Its origins in both cell death and damage and active excretion make it highly dynamic and responsive to immune, infectious, and inflammatory disease^6–8^. However, cfRNA suffers from low biomass and is susceptible to preanalytical factors that impact reproducibility. It is also relatively new to clinical research and thus has not yet been implemented extensively at the point of care^9^. Further evaluation is therefore needed to confirm its diagnostic performance relative to more established proteomic approaches.

To address these gaps, we investigated the agreement and diagnostic performance of high-throughput analysis of protein and RNA in plasma. Previous studies have compared protein and RNA profiles in other tissues^10,11^, but no direct comparisons of cfRNA and protein levels in plasma, where these analytes have high clinical relevance, have been performed. We used cfRNA sequencing and proteomics profiling by SomaScan for two overlapping pediatric inflammatory syndromes—KD and MIS-C. Our aims were ***i)*** to compare the abundance and correlation of cfRNA and proteins measured in matched plasma samples, ***ii)*** to identify both unique and shared molecular signatures associated with each condition and its subtypes, and ***iii)*** to evaluate the performance of cfRNA- and protein-based models in classifying KD and MIS-C. Although we observed no correlation between plasma RNA and protein levels as measured by RNA sequencing and SomaLogic proteomics, we found that highly accurate classifiers could nonetheless be developed to elucidate disease pathogenesis and heterogeneity based on both analytes. This study underscores the potential utility of differing molecular assays to refine diagnostic and therapeutic strategies.

## RESULTS

### Clinical Cohort

We enrolled and analyzed plasma samples from 263 pediatric patients diagnosed with Kawasaki Disease (KD) or Multisystem Inflammatory Syndrome in Children (MIS-C) at Rady Children’s Hospital San Diego (RCHSD) / University of California San Diego (UCSD). Of these, 155 samples were profiled by cfRNA sequencing (108 KD, 47 MIS-C) and 171 were analyzed using the SOMALogic SomaScan platform (70 KD, 101 MIS-C), with 63 samples profiled by both methods (**Fig. 1A**). KD cases were further stratified into clinical subtypes according to classification criteria defined by Wang et al.^12^. All samples were collected during the acute phase of illness and prior to treatment with intravenous immunoglobulin (IVIG).

**Fig. 1.**
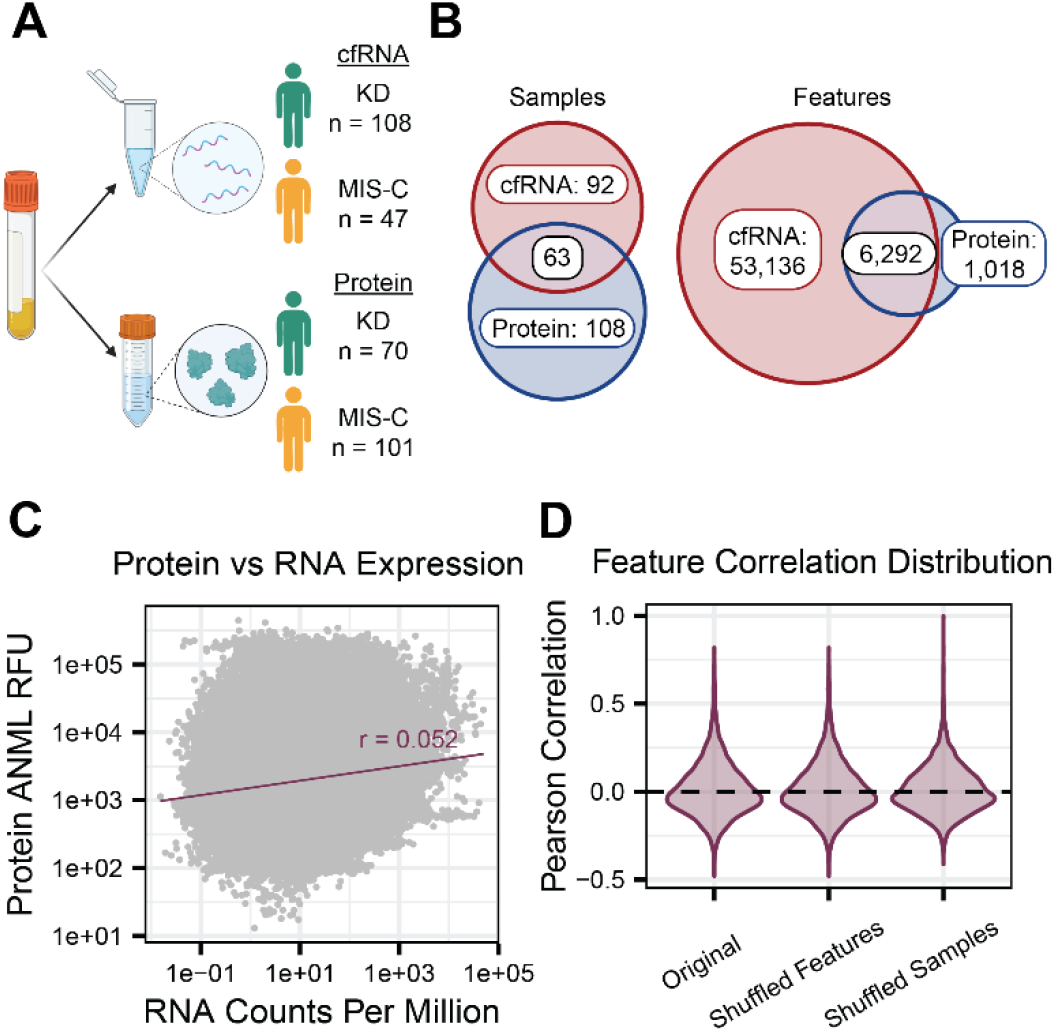
Overview of study analytes and samples. **(A)** cfRNA sequencing and SomaScan from plasma sample composition by condition and analyte. **(B)** Sample and feature overlap across cfRNA and protein assays. **(C)** Correlation of feature expression in 63 matched samples. **(D)** Distributions of individual feature correlations compared to correlations of randomized samples and features.

### Lack of correlation of cfRNA and protein levels

To assess how plasma cfRNA and proteins compare as potential diagnostic biomarkers, we first compared the biomolecular features captured by each assay. We conducted untargeted cfRNA profiling by RNA sequencing and assigned the reads to a total of 59,428 annotated features. For proteomics, we used the SOMALogic SomaScan affinity-based platform, which contains 7,310 targeted aptamers to 6,397 unique proteins^3^. Of these, 6,292 overlapped with the cfRNA features (**Fig. 1B**). We evaluated global similarities by measuring the correlation of all molecules in paired samples. We observed no overall correlation (Pearson r=0.052, Spearman rho = 0.131) when comparing counts per million (cfRNA) to adaptive normalization by maximum likelihood relative fluorescence units (ANML RFUs, protein, **Fig. 1C**). At the individual feature level, protein and RNA levels were uncorrelated (median Pearson r=-0.008), suggesting that cfRNA and protein capture largely distinct plasma biomolecular landscapes. Randomly shuffling sample identifiers and feature labels produced similar distributions centered around zero correlation (**Fig. 1D**).

### Differentially abundant features and pathways

Given the observed lack of correlation between cfRNA and proteins in plasma, we asked how each analyte distinguished between KD and MIS-C—two clinically overlapping yet pathologically distinct pediatric inflammatory conditions^13^. We performed differential abundance and pathway analyses for both cfRNA and proteins and compared their significant features (**Methods**). Overall, we identified 439 differentially abundant cfRNA transcripts and 63 differentially abundant proteins (BH adjusted p-value < 0.05 and |log2 fold change| > 1, **Fig. 2A**).

**Fig. 2.**
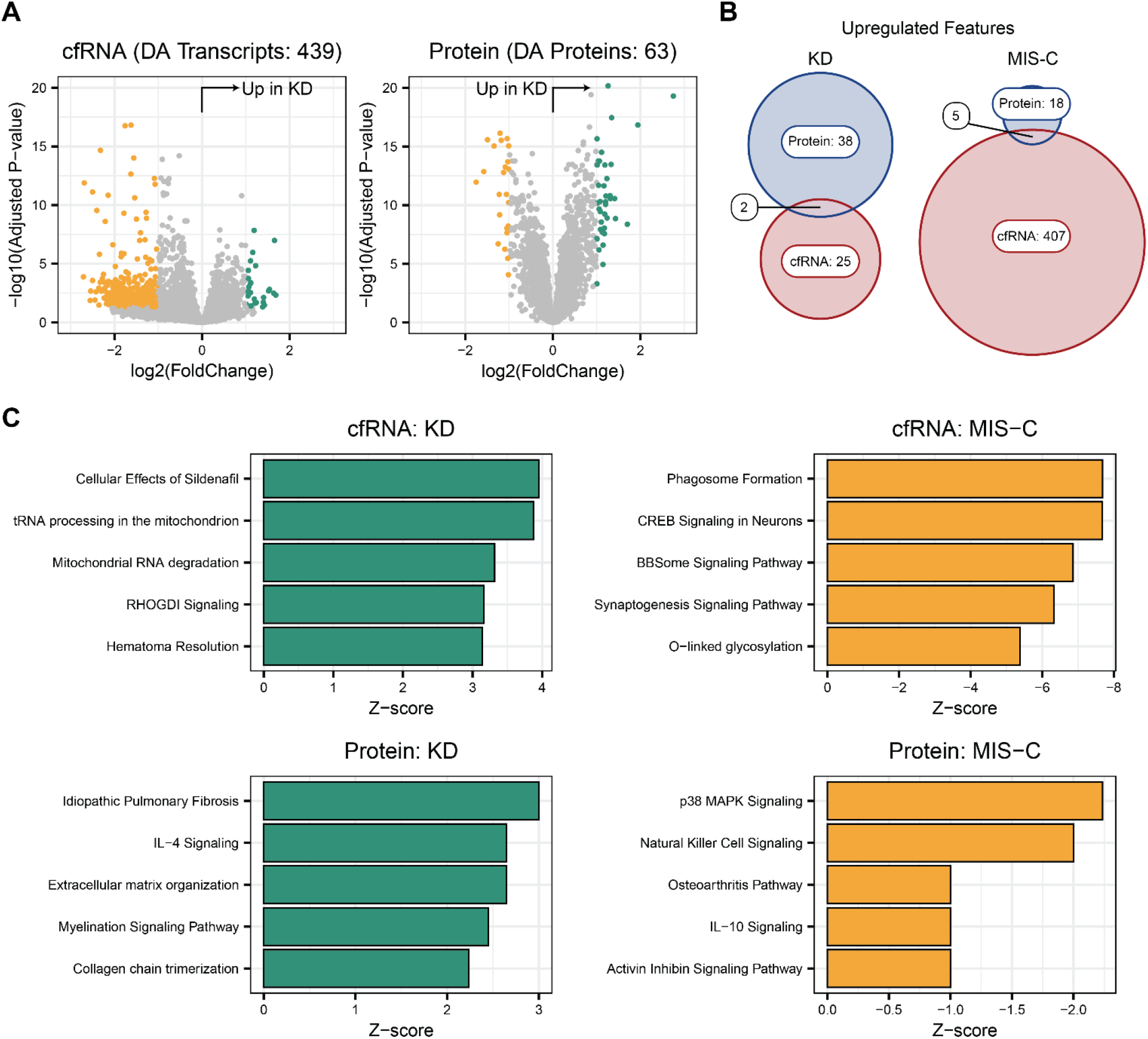
Comparisons in differential abundance and Ingenuity Pathway Analysis (IPA). **(A)** Volcano plot of differentially abundant features found in cfRNA and protein. **(B)** Overlap in differentially abundant features by condition found in cfRNA and protein. **(C)** Top 5 upregulated pathways found in IPA for each condition and analyte.

While cfRNA and proteins showed similar ranges of fold changes (log2: cfRNA = [-2.71, 1.68], protein = [-1.75, 2.75]), 68 cfRNA features—versus only one protein—exhibited a log2 fold change greater than 2 (**Fig. 2A**). We observed more differentially abundant proteins upregulated in KD than cfRNA (protein n = 40, RNA n = 27, log fold change > 0), with only two features, COL10A1 and CCL5, upregulated across both datasets. The differences between cfRNA and protein became more pronounced in MIS-C, where 412 features were upregulated in cfRNA but only 23 in protein (**Fig. 2B**). Among these, 5 (*IL1R1, LAP3, CXCL10, CNDP1, VSIG4*) were upregulated in both analytes. Several of these (IL1R1, CXCL10, VSIG4) are known to contribute to immune activation and response^14–16^,suggesting that despite limited overall overlap at the molecule level, cfRNA and proteins may converge on immune-related features.

To contextualize the differentially abundant cfRNA and proteins within larger-scale cellular functions, we performed pathway analyses using Ingenuity Pathway Analysis (IPA) (**Methods, Fig. 2C**). In KD, the top enriched pathways in cfRNA highlighted active immune and inflammatory responses, most notably hematoma resolution and cellular effects of Sildenafil^17^, suggesting transcriptional regulation of acute vascular pathology. In contrast, protein-based pathways in KD included IL-4 signaling, myelination signaling, and collagen chain trimerization, which reflect downstream effects on inflammation, cellular function, and tissue remodeling. In MIS-C, cfRNA analysis revealed disruptions in neuronal and intracellular signaling pathways (e.g. phagosome formation and synaptogenesis). Conversely, protein analysis pointed to inflammatory and metabolic dysregulation, such as p38 MAPK signaling, natural killer cell signaling, and IL-10 signaling, highlighting active immune processes not captured at the transcript level. These findings suggest that cfRNA and protein analyses provide complementary insight, with cfRNA capturing upstream regulatory processes and protein revealing downstream functionality. This analysis reinforces the complementary nature of both analytes to gain a more complete understanding of pathophysiology.

### Both cfRNA and Protein Can Classify KD and MIS-C

Because no molecular tests currently exist for KD or MIS-C, we evaluated whether plasma cfRNA, proteins, or both can be used to develop diagnostic signatures. To do so, we trained machine learning classifiers using repeated cross-validation (**Methods**). Specifically, we trained a generalized linear model with LASSO regression (GLMNETLasso) to optimize biomarker panel size via regularization. Across 250 iterations, we observed similar performance for models trained on cfRNA and protein profiles, with a median test AUC across iterations of 0.940 for cfRNA and 0.937 for protein (p = 0.8602, **Fig. 3C**). While performance was similar for both analytes, the cfRNA classifiers achieved this high performance using fewer features (**Fig. 3D**). To dissect whether these outcomes were influenced by sample composition or the number of available features, we repeated the cross-validation analysis for three scenarios: matched samples (n = 63), matched features (n = 6,292), and matched samples and features (**Fig. 3E**). Matching samples reduced performance for both cfRNA (median AUC = 0.896) and protein (median AUC = 0.870), highlighting the impact of sample size on classification accuracy (p = 6.528e-06). When only features were matched, proteomics (median AUC = 0.933) outperformed cfRNA (median AUC = 0.877, p = 2.2e-16), suggesting that the broader biomarker repertoire of cfRNA is a key driver of its classification power. With both samples and features matched, cfRNA (median AUC = 0.883) and protein (median AUC = 0.857) performed similarly with a slight improvement in cfRNA (p = 3.148e-4), again indicating comparable performance in distinguishing KD from MIS-C. Finally, we examined which biomarkers were most frequently selected across the full dataset and target space. Six transcripts (RN7SL752P, TCF7, CA1, IFIT2, GBP5, MAP3K7CL) appeared in more than 50% of cfRNA-based models, while seven proteins (COL10A1, PLA2GA, DEFB4A, CCL17, PI3, LAP3, PDIA5) were selected in the majority of protein-based models. **Fig 3F**) Notably, LAP3 was one of the most frequent features selected in both analytes, which suggests a more ubiquitous expression across RNA and protein that is relevant to KD.

**Fig. 3.**
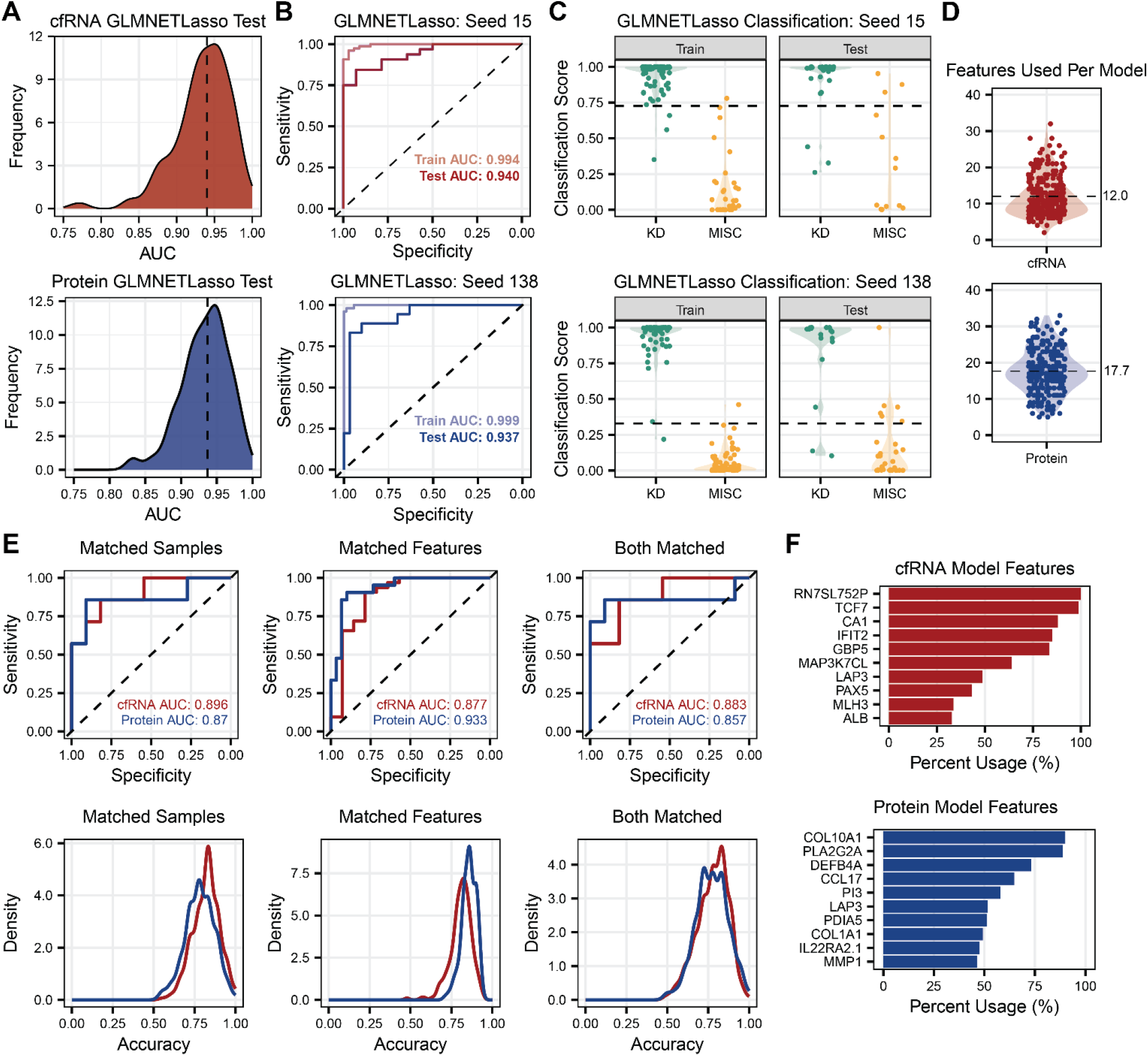
Machine learning classifiers constructed using a GLMNET model with Lasso regression. **(A)** Distribution of AUC values across 250 iterations for cfRNA- and protein-based classifiers. **(B)** AUC curves and classification results for train and test splits for the seed with the median AUC (cfRNA seed 15, protein seed 138). **(C)** Classification results of the median-performing seed for cfRNA and protein. **(D)** Distribution of total features used in the final model across 250 iterations for cfRNA and protein. **(E)** AUC curves of the median AUC seed and test accuracy distribution for cfRNA and protein when using only matched samples, matched features, and both matched samples and features. **(F)** Top 10 cfRNA and protein features by usage in total percentage of final models.

### KD subtyping based on cfRNA and protein profiles

A recent study demonstrated that KD patients can be stratified into four subtypes based on 14 clinical variables, each associated with distinct clinical outcomes and molecular profiles^12^. These subtypes align with specific hallmark features: subtype 1 is linked to elevated liver involvement, subtype 2 to increased band neutrophil activity, subtype 3 to pronounced lymphadenopathy, and subtype 4 to a lower average age of onset of disease. Given the different molecular perspectives offered by cfRNA and protein, we examined whether these more granular phenotypic differences were more clearly captured by one analyte compared to the other. We performed differential abundance analyses of cfRNA and proteins for each KD subtype relative to MIS-C. All KD subtypes exhibited both differentially abundant cfRNA transcripts and proteins, except subtype 2, in which only one protein, CCL17, was differentially abundant when compared to MIS-C (**Fig. 4A**). This suggests that subtype 2 may share greater molecular similarity with MIS-C than with other KD subtypes. Consistent with these findings, repeated cross-validation analyses revealed comparably high accuracy for cfRNA- and protein-based models in distinguishing the other KD subtypes from MIS-C; however, classification accuracy for subtype 2 was notably lower in the protein-based models at only 52%, suggesting the model is incapable of distinguishing subtype 2 from MIS-C (**Fig. 4B**). Together, these results support an elevated molecular similarity between subtype 2 and MIS-C that may influence clinical presentation.

**Fig. 4.**
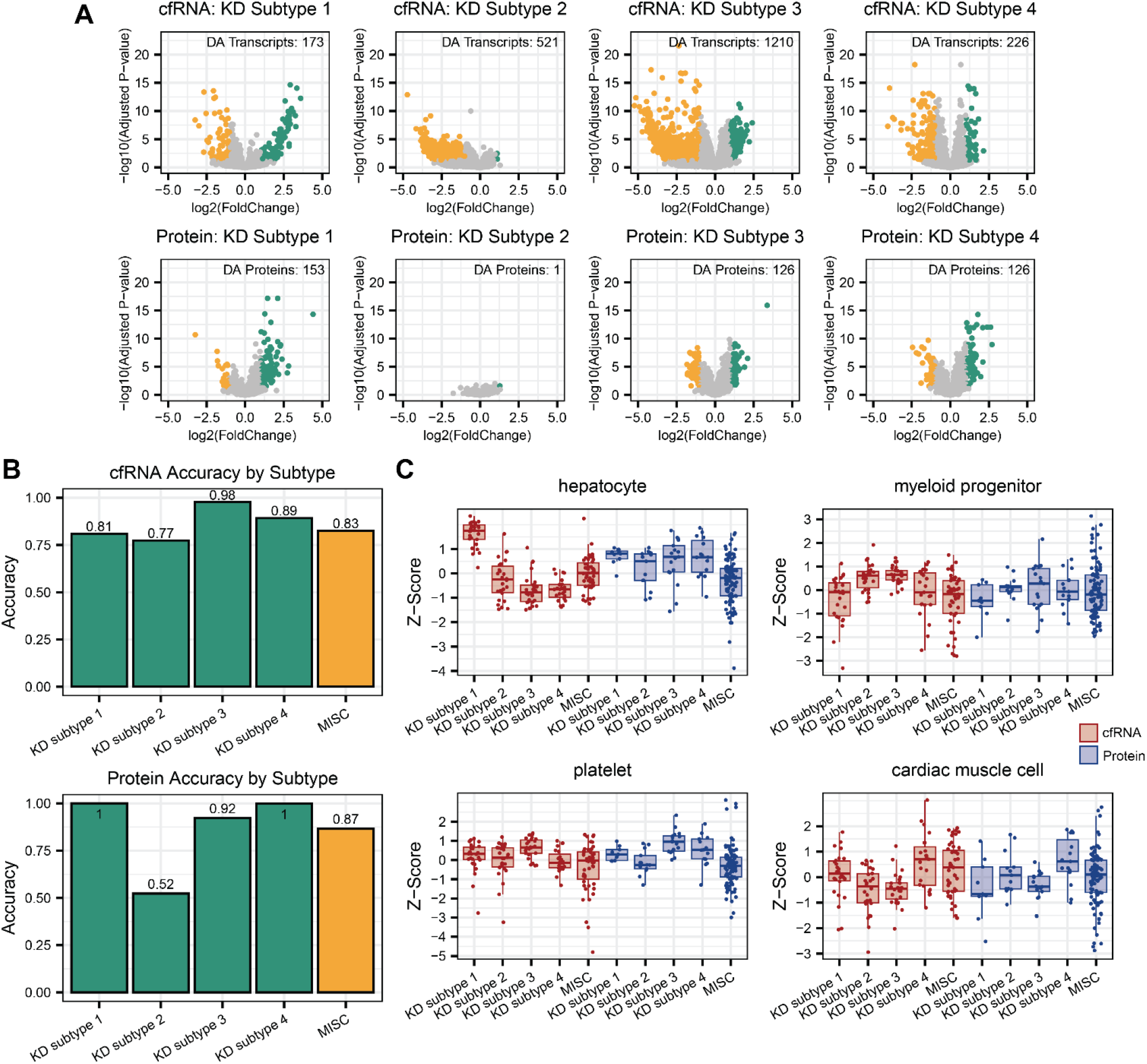
Comparative analysis of KD subtypes using cfRNA and protein. **(A)** Volcano plots for each subtype versus MIS-C in both cfRNA and protein. **(B)** Rate of correct classification of each subtype and MIS-C across 250 iterations of machine learning classifiers. **(C)** Boxplot of differential cell types for each KD subtype by Z-score using SSGSEA.

To further explore subtype-specific differences, we performed single-sample gene set enrichment analysis (ssGSEA) using the Tabula Sapiens cell type reference (**Methods, Fig. 4C**). The top enriched cell type differed across KD subtypes and often diverged between cfRNA and protein data. For instance, subtype 1—known for liver-enriched pathophysiology—exhibited increased liver-associated transcripts in cfRNA, yet a similar pattern was not seen in the protein data, despite hepatocytes being the most enriched cell type in the protein data. In contrast, immune cell types—particularly granulocytes in subtype 2 and CD4-positive helper T cells in subtype 4—displayed more consistent enrichment patterns across both cfRNA and protein, underscoring a shared underlying immune activation. These results suggest that cfRNA may better capture the subtype-specific clinical phenotypes, but better concordance between cfRNA and protein may be in detecting immune-related molecular features and pathways.

## DISCUSSION

We compared two high-throughput platforms for plasma bio-analyte analysis, RNA sequencing of cell-free RNA (cfRNA) and SomaScan proteomics of circulating proteins, to evaluate both their concordance and their utility in disease classification. Previous studies of mRNA–protein relationships in tissues and whole blood have reported only modest correlations^10,11^ (0.3–0.6), reflecting the multiple biological mechanisms that decouple transcript and protein levels. Post-transcriptional regulation, protein turnover, and dynamic transitions from transcription to translation^18–21^ may all contribute to this discordance in addition to technical differences in the measurement platforms.

Here, we found that cfRNA transcripts and proteins are essentially uncorrelated on a feature-by-feature basis, suggesting that each platform captures a distinct aspect of underlying biology. Indeed, the correlations between plasma RNA and proteins observed here are considerably weaker than the correlations reported in tissues and whole blood, indicating that additional factors such as release mechanisms, stability, and clearance may further disrupt any residual tissue-level concordance. Notably, in MIS-C, there were substantially more differentially abundant transcripts than proteins, which could reflect increased cfRNA release triggered by tissue damage during systemic inflammation. Despite minimal overlap at the individual molecular level, both cfRNA and protein converged on immune pathways, with cfRNA often revealing upstream regulators (e.g., interferon-inducible genes) and protein data highlighting downstream effectors (e.g., inflammatory mediators). Together, these findings suggest that combining cfRNA and protein measurements can lead to a more comprehensive perspective on pathophysiological processes.

Previous work has established the ability of cfRNA to correctly classify KD and MIS-C^6^. When we used machine-learning models to classify KD vs. MIS-C in both cfRNA and protein, both classifiers achieved high accuracy but selected different sets of diagnostic features. The cfRNA models emphasized immune-related transcripts (e.g. *TCF7*, which regulates the development of immune cells and inflammation^22^, *IFIT2*, which regulates interferon expression^23^, *GBP5*, which contributes to inflammasome assembly and is induced by interferon^24^, and *MAP3K7CL*, which has been implicated in pediatric sepsis^25^. Of note, the models all included *RN7SL752P*, a pseudogene, suggesting additional diagnostic information contained within non-protein-coding transcripts. Protein models also chose immune-related features (e.g. DEFB4A, a defensin that can be and found in vascular endothelial cells^26^, COL10A1 and COL1A1, involved in immune cells recruitment^27,28^, PI3, an antimicrobial peptide^29^, and LAP3, a metallopeptidase implicated in T cell depletion^30^. CCL17, a cytokine implicated in immune processes including arthritic disease and skin inflammation^31,32^, was the most differentially abundant protein we observed in KD, suggesting a possible connection between its expression and the clinical presentation of skin inflammation seen in KD patients.

Further analysis of KD subtypes revealed that subtype 2 proved the most challenging to distinguish from MIS-C, with few or no differentially abundant proteins identified. This may stem from an infectious trigger common to both conditions, as suggested by a higher incidence of KD shock syndrome (KDSS) in subtype 2. A final point of interest was the discrepancy in liver-derived signals for KD subtype 1: cfRNA revealed elevated liver transcripts, yet proteomic data did not mirror this trend, even though classical protein-based liver function tests align with clinical pathology. This suggests that cfRNA more specifically reveals tissue injury as compared to proteomic data. These findings illustrate how clinical assays, affinity-based proteomics, and cfRNA profiling each capture unique facets of disease.

High-throughput proteomics platforms continue to evolve, with debate ongoing over their optimal role in large-scale research and clinical settings. While mass spectrometry has been a gold standard for protein discovery, affinity-based methods such as SomaScan and Olink can profile thousands of targets from minimal volumes but are susceptible to platform biases and remain expensive ($750-1000). Proteomic platforms have shown lack of interplatform agreement, coverage, and reproducibility^1^, underscoring the importance of platform choice in interpreting results^33^. In contrast, cfRNA profiling offers a real-time snapshot of gene expression at comparatively lower costs (∼$150 per sample) and the option to convert detection of cfRNA to a quantitative PCR platform. However, widespread clinical implementation of cfRNA assays will require new protocols, workflows, and infrastructure.

Taken together, our results underscore the complementary nature of cfRNA and protein as blood-based biomarkers. Ultimately, integrating cfRNA and protein analyses together could enhance molecular characterization, improve disease classification, and guide more targeted therapeutic approaches.

## METHODS

### Ethics Statement

The Cornell University IRB for Human Participants (2012010003), New York, NY, approved the protocols for this study. All samples and patient information were deidentified for analysis and shared with collaborating institutions. At UCSD, the IRB reviewed and approved collection and sharing of samples and data (IRB #140220). Signed consent and assent as appropriate were provided by the parent(s) and pediatric patient, respectively.

### Clinical cohort

We examined a cohort of 263 pediatric patients diagnosed with KD or MIS-C at Rady Children’s Hospital–San Diego between 2006 and 2021. Two experienced KD clinicians (AHT and JCB) confirmed each diagnosis in accordance with American Heart Association criteria for both complete and incomplete KD (28356445), and clinical data were prospectively recorded at the time of diagnosis. Among these patients, plasma samples for 155 (n=108 KD and n=47 MIS-C) were analyzed by cfRNA sequencing. In a previous study (PMID: 37598693), KD cases were further categorized into four subgroups (n=26 Subgroup 1, n=26 Subgroup 2, n=26 Subgroup 3, n=23 Subgroup 4). Additionally, we performed SomaScan proteomic analysis on samples from 171 patients (70 KD and 101 MIS-C), which included 54 KD patients who had been categorized by subgroups (n=9 Subgroup 1, n=13 Subgroup 2, n=17 Subgroup 3, n=15 Subgroup 4). A total of 63 patient samples were included in both the cfRNA and proteomic analysis.

### Sample collection and cell-free RNA sequencing

Plasma samples were collected at UCSD in lavender-top EDTA tubes prior to treatment. Samples were centrifuged at 2,000 g for 10 minutes and stored at -80°C prior to shipping. cfRNA was extracted and cDNA libraries were prepared for sequencing. Sequencing data was analyzed as previously described (39240972).

### Sample Quality Control

Quality control was performed using the sequencing data by assessing DNA contamination, rRNA contamination, total counts, and RNA degradation. DNA contamination was estimated based on the intron-to-exon read mapping ratio, while rRNA contamination was measured with SAMtools (v1.14). Total counts were computed using featureCounts30 (v2.0.0), and degradation was estimated by calculating the 5′-to-3′ read alignment bias with Qualimap31 (v2.2.1). Samples were excluded if the intron-to-exon ratio exceeded 3, if total counts were below 75,000, or if the 5′-to-3′ bias exceeded 2.

### SomaScan analysis

Plasma samples from the same collection used for cfRNA were sent to SomaLogic for proteomic profiling using the SomaScan 7K assay. Raw and ANML SMP normalized data using a pediatric normalization reference were provided by SomaLogic.

### Differential abundance analysis

We assessed transcript abundance using a negative binomial model and applied variance stabilizing transformation (VST) via the DESeq2 R package (25516281).

### Machine learning

The data were split into training and test sets at a 70:30 ratio over 250 iterations to minimize random seed variability. Differential abundance analysis was then performed on the training data, filters by adjusted P-value < 0.05, base mean abundance > 50, and an absolute log2 fold change > 1. The test set was excluded from threshold determination to prevent overfitting.

Raw counts for the training, validation, and test sets were each normalized via variance-stabilizing transformation (VST), and transcript subsets were formed based on the selected features. Variance stabilization was performed using DESeq2, with the dispersion function from the training set applied to transform the validation and test sets. Model training incorporated fivefold cross-validation and grid search hyperparameter tuning using GLMNET with Lasso regularization, which reduced irrelevant feature coefficients to zero and selecting only the most informative transcripts.

For each model, classification score thresholds were established using Youden’s index on the training data. The trained models were subsequently used to predict labels in the validation set, and performance was measured by the area under the ROC-AUC. Predictions on the test set employed the same Youden’s index threshold derived from the training set.

### Cell type enrichment analysis

Cell enrichment analysis was performed via single sample gene set enrichment analysis (ssGSEA) using the GSVA package in R^34^ (ssgsea.norm=FALSE, tau=0.75, parallel.sz=10, min.sz=10), instead using Tabula Sapiens v2 as the reference gene set to annotate enrichments to cells rather than pathways. We performed pathway and network analysis using Ingenuity Pathway Analysis^35^ (IPA, QIAGEN).

### Ingenuity Pathway Analysis

We performed pathway and network analysis using Ingenuity Pathway Analysis (IPA, QIAGEN) separately on cfRNA and protein expression datasets. The top five enriched pathways were determined by successive removal of pathways sharing greater than 75% of annotated features with the top enriched pathway. Pathways annotated as cancer-specific were excluded from the results to avoid overrepresentation.

### Statistical analyses

All statistical analyses were conducted using R (v4.1.0). We determined statistical significance with two-sided Wilcoxon signed-rank tests and Mann-Whitney U tests. In boxplots, the box edges represent the 25th and 75th percentiles, the central line indicates the median, and whiskers extend to 1.5 times the interquartile range beyond the hinge.

## Data availability

De-identified RNA-seq count matrices and have been deposited in the NCBI Gene Expression Omnibus (GEO) database (GSE255555) and will be made publicly available upon publication. All associated code has been uploaded to GitHub and will also be accessible upon publication (https://github.com/adb258/cfRNA_proteins_manuscript.git).

## AUTHOR CONTRIBUTIONS

A.B., C.J.L., J.C.B., and I.D.V. designed research; A.B., C.J.L., J.L., E.B. processed samples; A.B., C.J.L., J.K., C.S., J.L., E.B., A.H.T., J.C.B., C.Y.C., and I.D.V. analyzed data; C.S., A.H.T., and J.C.B. identified and collected patient samples and clinical metadata; and A.B., C.J.L., and I.D.V. wrote the paper.

## ACKNOWLEDGEMENTS

We thank the patients and their families for their help to further our understanding of pediatric inflammatory conditions. We thank the Cornell Genomics Center for help with sequencing libraries. We thank the members of the PEMKDRG: Lukas Austin-Page, MD, Amy Bryl, MD, Joelle Donofrio-Ödmann, MD, Atim Ekpenyong, MD, David Gutglass, MD, Scott Herskovitz, MD, Paul Ishimine, MD, John Kanegaye, MD, Margaret Nguyen, MD, Mylinh Nguyen, MD, Kristy Schwartz, MD, Stacey Ulrich, MD, Tatyana Vayngortin, MD, and Elise Zimmerman, MD. This work was funded by NIH/National Institute of Child Health and Human Development grants R61HD105618, R33HD105618, and R33HD105593 (C.S., A.H.T., J.C.B., and I.D.V.) and the Macklin foundation. The funders had no role in study design, data collection and analysis, decision to publish, or preparation of the manuscript.

