## Supplemental Information for "Minimal Correlation but Complementary Diagnostic Utility for Plasma Cell-free RNA and Proteins"

**Supplementary Table 1.**
Patient cohort demographic information.

| | | ***Cell-free RNA*** | | ***Protein*** | |
|---|---|---|---|---|---|
| Variables | **Group** | **KD** | **MIS-C** | **KD** | **MIS-C** |
| n | | 108 | 47 | 70 | 101 |
| KD Subtype, n (%) | Subtype 1 | 26 (24.07) | | 9 (18.57) | |
| | Subtype 2 | 26 (24.07) | | 13 (18.57) | |
| | Subtype 3 | 26 (24.07) | | 17 (24.29) | |
| | Subtype 4 | 23 (21.30) | | 15 (21.43) | |
| | Unknown | 7 (6.48) | | 16 (22.86) | |
| Sex, n (%) | Male | 61 (56.48) | 35 (74.47) | 45 (64.29) | 66 (65.35) |
| | Female | 47 (43.52) | 12 (25.53) | 25 (35.71) | 35 (34.65) |
| Age, mean | | 4.05 | 9.09 | 4.05 | 8.70 |
| Race, n (%) | Asian | 19 (17.59) | 2 (4.26) | 12 (17.14) | 4 (3.96) |
| | Black/African American | 4 (3.70) | 8 (17.02) | 2 (2.86) | 17 (16.83) |
| | Caucasian | 20 (18.52) | 5 (10.64) | 19 (27.14) | 8 (7.92) |
| | Hispanic | 33 (30.56) | 27 (57.45) | 26 (37.14) | 55 (54.46) |
| | More than one race | 28 (25.93) | 4 (8.51) | 11 (15.71) | 13 (12.87) |
| | Other | 4 (3.70) | 1 (2.13) | 0 (0.00) | 2 (1.98) |
| | Unknown | 0 (0.00) | 0 (0.00) | 0 (0.00) | 2 (1.98) |
| Matched in both analytes, n (%) | | 24 (22.22) | 39 (76.60) | 24 (34.29) | 39 (38.61) |

**Data S1. (separate file)**
DESeq2 results from KD vs MIS-C comparisons in cell-free RNA.

**Data S2. (separate file)**
DESeq2 results from KD vs MIS-C comparisons in proteins.

**Data S3. (separate file)**
IPA results from cell-free RNA transcript differential abundance analysis from KD vs MIS-C comparisons.

**Data S4. (separate file)**
IPA results from protein differential abundance analysis from KD vs MIS-C comparisons.
